# Potential of genomic selection for improvement of resistance to Ostreid Herpes virus in Pacific oyster (*Crassostrea gigas*)

**DOI:** 10.1101/754473

**Authors:** Alejandro P. Gutierrez, Jane Symonds, Nick King, Konstanze Steiner, Tim P. Bean, Ross D. Houston

## Abstract

In genomic selection (GS), genome-wide SNP markers are used to generate genomic estimated breeding values (gEBVs) for selection candidates. The application of GS in shellfish looks promising and has the potential to help in dealing with one of the main issues currently affecting Pacific oyster production worldwide, which is the “summer mortality syndrome”. This causes periodic mass mortality in farms worldwide and has mainly been attributed to a specific variant of the Ostreid herpesvirus (OsHV-1-μvar). In the current study, we evaluated the potential of genomic selection for host resistance OsHV in Pacific oysters, and compared it to pedigree-based approaches. An OsHV-1 disease challenge was performed using an immersion-based virus exposure treatment for oysters for seven days. 768 samples were genotyped using the medium density SNP array for oysters. GWAS was performed for the survival trait using a GBLUP approach in BLUPF90 software. Heritability ranged from 0.25±0.05 to 0.37±0.05 (mean±s.e) based on pedigree and genomic information, respectively. Genomic prediction was more accurate than pedigree prediction, and SNP density reduction had little impact on prediction accuracy until marker densities dropped below ∼500 SNPs. This demonstrates the potential for GS in Pacific oyster breeding programs and importantly, demonstrates that a low number of SNPs might suffice to obtain accurate gEBVs, thus potentially making the implementation of GS more cost effective.

## Introduction

The use of genomic information to predict breeding values for selection candidates has become commonplace in advanced breeding programmes. Genomic selection (GS, proposed by Meuwissen *et al.* (2001), uses genome-wide markers to capture genetic variation in the trait of interest, even if the trait is highly polygenic. GS involves measurements of trait values and genotypes in a reference or training population, training of the genomic prediction model, and then use of this model to predict genomic breeding values (gEBVs) for selection candidates (Goddard and Hayes 2007).

High throughput genome-wide genotyping is a major component of genomic selection programmes. SNP arrays have enabled routine genotyping, facilitating the typing of many thousands of SNP markers dispersed throughout the genome of multiple individuals of the target species. Accordingly, SNP arrays have been developed for many important finfish aquaculture species such as Atlantic salmon, rainbow trout, catfish and carp among others (Houston *et al.* 2014; Yáñez *et al.* 2016; Palti *et al.* 2015; Liu *et al.* 2014; Xu *et al.* 2014). In addition, two SNP arrays have been recently developed for Pacific oyster (*C. gigas*); a combined-species medium density array for Pacific oyster and European flat oyster (*O.edulis)* (Gutierrez *et al.* 2017) and a high density array for Pacific oyster (Qi *et al.* 2017). Moreover, a high density linkage map containing ∼20K SNPs has recently been created and aligned with the physical reference genome assembly (Gutierrez *et al.* 2018a; Zhang *et al.* 2012). Using such arrays, several studies have demonstrated that genomic selection for aquaculture species results in improved accuracy compared to traditional pedigree-based approaches; for example in Atlantic salmon (Robledo *et al.* 2018), coho salmon (Barría *et al.* 2018), rainbow trout (Vallejo *et al.* 2018), common carp (Palaiokostas *et al.* 2018b), and Pacific oyster (Gutierrez *et al.* 2018b).

One of the main issues currently affecting oyster production worldwide is the “summer mortality syndrome”. These events cause periodic mass mortality in farms worldwide and have been mainly attributed to a specific variant of the ostreid herpesvirus (OsHV-1-μvar) (Segarra *et al.* 2010), amongst other factors (de Lorgeril *et al.* 2018; Petton *et al.* 2015; Malham *et al.* 2009). Selective breeding to improve resistance to OsHV-1 may have potential as a prevention strategy, and there is significant additive genetic variation in survival during OsHV-1 infection, with estimated heritability values ranging from 0.12 to 0.63 (Azéma *et al.* 2017; Camara *et al.* 2017; Dégremont *et al.* 2015b; Gutierrez *et al.* 2018a). However, QTL and GWAS approaches to investigate the genetic architecture of host resistance to the virus have suggested that the trait may be polygenic (Gutierrez *et al.* 2018a; Sauvage *et al.* 2010). For that reason, marker-assisted selection for OsHV-1 resistance is unlikely to be effective, and the trait is a good candidate for genomic selection.

Substantial efforts have been made to establish selective breeding programs for Pacific oyster, with OsHV-1 resistance as a primary target trait (Dégremont *et al.* 2010; Dégremont *et al.* 2015a; Camara and Symonds 2014). An encouraging response to selection for resistance has been observed in Pacific oyster spat after four generations of mass selection (Dégremont *et al.* 2015c), while family based selection breeding programs have shown encouraging results after a few generations (Camara *et al.* 2017). Given that Pacific oysters account for 98% of global oyster production, which was estimated at ∼0.6 M tons in 2016 (FAO, 2018), this pathogen is a significant problem for global aquaculture. Genomic tools can enhance selective breeding in aquaculture species via improvements in selection accuracy corresponding to increased genetic gain, together with improved control of inbreeding (Houston 2017).

The aim of this study was to investigate the genetic architecture of resistance to OsHV-1 infection in a Pacific oyster population from New Zealand by a disease challenge experiment followed by a genome wide association study (GWAS), and to evaluate the use of genomic prediction to inform the implementation of genomic selection in selective breeding programmes.

## Methods

### Source of oysters and disease challenge

Families were produced at the Cawthron Institute’s hatchery in Nelson, NZ as part of the 2015 cohort in an ongoing commercial selective breeding program. Families within this cohort can be separated in two groups: the “Nucleus” group which originated from survivors of OsHV-1 exposure with expected higher levels of resistance to OsHV-1; and the “Down-selected” group originating from families that have shown poor survival to the virus. Estimated breeding values based on field challenge survival were used to select the parents from the previous 2013 cohort to form the nucleus and down-selected full-sib families.

The OsHV-1 challenge experiment was performed on 1860 animals approximately 4 months post-spawning using the immersion challenge model described in Camara et al. (2017). In brief, viral stock is obtained from oysters with confirmed high virus loads which are homogenised, and after serial filtering steps the supernatant is cryopreserved in a 10 % glycerol and 10 % foetal calf serum solution. The same process was used to prepare a negative control stock from virus-free oysters.

Subsequently, groups of 20 oysters derived from each of 31 full sibling families were exposed to one of three treatments; (i) a high concentration of the virus, (ii) a low concentration of the virus, and (iii) a mock-challenged negative control (total n = 1860). Inoculation for the high virus concentration (hv) was prepared by diluting 4 mL of the virus stock in 7.7 L of artificial seawater (ASW), same procedure was followed for the negative control (no virus stock). The inoculation for the low virus concentration (lv) was a 10-fold dilution with ASW of the high virus concentration water. All families from the 2015 cohort with sufficient available spat were challenged. The oysters were randomly sampled from their family-specific upwelling tanks in the virus-free nursery, cleaned and placed in family-specific mesh bags which were transferred to three 120 L tanks in a pre-determined pattern to spread the families evenly throughout the tank in a grid system. Inoculation water (7 L/tank) was added to the tanks, and after 16 h incubation a further 73 L ASW was added to each tank with constant aeriation and temperature of 21 °C. Water samples (100 µL) were collected daily from each tank and frozen prior to qPCR analysis to confirm viral load in the treatments and control. ASW was replaced every second day with the same volume of newly prepared ASW and microalgal feed in the form of 120 ml axenically-cultured Isochrysis galbana was added to each tank. The oysters were assessed for signs of life daily for 7 days after inoculation. Dead oysters were removed, counted, and frozen until DNA extraction.

### SNP array genotyping

Genomic DNA was extracted from the whole oyster (minus the shell) using the E.Z.N.A. Mollusc DNA Kit (Omega Biotek), quantified on Qubit and the DNA integrity was checked on a 1% agarose gel. Following quality control (QC) and considering the 384 well configuration of the array, 768 samples (718 progenies and 44 parents) were sent for genotyping to ThermoFisher Scientific (Santa Clara, USA) using the recently developed Affymetrix SNP medium density SNP array for oysters (Gutierrez *et al.* 2017). All dead oysters were genotyped, as well as 16 surviving oysters from each family (8 from each treatment) when possible, therefore, the number of genotyped individuals per family ranged from 16 to 40 depending on the DNA quality and the number of mortalities per family in each condition (hv or lv) (Table S1 and S2). After genotyping, initial QC steps were performed using the Axiom Analysis Suite v2.0.0.35, and 762 individuals and 22,535 SNPs were retained following the “best practices workflow”, which included ‘PolyHighResolution’ and ‘NoMinorHom’ SNPs only, a sample and SNP call threshold of 90 %, plus a manual inspection and removal of selected SNPs with unusual clustering patterns. The SNPs were then checked for Mendelian errors using Plink 1.9 (Chang *et al.* 2015), leaving a total of 21,338 SNPs and 762 individual animals that were used for the linkage map construction. Final filtering of the SNP set was performed using the GenABEL package (Aulchenko *et al.* 2007) in R, using the *check.markers* module to retain SNPs with a MAF > 0.01 and allow a deviation from Hardy-Weinberg Equilibrium < 1 × 10^−6^, leaving 17,919 filtered SNPs that were used for heritability, GWAS and GS analyses.

### Linkage mapping

Due to the lack of a chromosome-anchored reference genome assembly for Pacific oyster, the genotype data were used to construct a high density linkage map. The maps from a total of 21,338 SNPs were constructed using the software Lep-map 3 (Rastas 2017). The nuclear families used for the generation of this map were assigned using the SNP data and the software Cervus (Kalinowski *et al.* 2007) as described by Gutierrez *et al.* (2017), and confirmed through the IBD module in Lep-map3. Putative erroneous or missing parental genotypes were re-called using the “ParentCall2” module. Linkage groups were identified using the “SeparateChromosomes2” module using a LodLimit = 31 and distortionLod = 1. Data were then filtered to remove markers from families showing significant segregation distortions (“dataTolerance=0.001”) and the “OrderMarkers2” module was applied to order the markers within the linkage groups. Individuals showing excessive recombination were also removed from the data as this indicated a potential problem with genotyping or family assignment for this individual. Additionally, markers that could not be assigned were positioned according to our previously developed linkage map (Gutierrez *et al.* 2018a).

### Estimation of genetic parameters for OsHV-1 resistance

Genetic parameters for the OsHV-1 resistance traits were estimated using a linear mixed model approach fitting animal as a random effect and tank as fixed effect, using ASReml 4 (Gilmour et al. 2014) with the following model, but adjusted to the logit-link function:

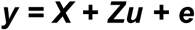

where **y** is the observed trait, **u** is the vector of additive genetic effects, **e** is the residual error, and **X** and **Z** the corresponding incidence matrices for fixed effects and additive effects, respectively. The (co)variance structure for the genetic effect was calculated either using pedigree (**A**) or genomic (**G**) matrices (i.e. **u** ∼ N(0, Aσ_a_ ^2^) or N(0, Gσ_a_ ^2^)), where G is the genomic relationship matrix and σ^2^ is the genetic variance. The heritability of the traits was estimated using the additive genetic variance and total phenotypic variance, as follows:

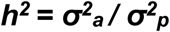

where σ^2^ _a_ is the additive genetic variance and σ^2^ _p_ is the total phenotypic variance which is a sum of *σ* ^*2*^ _*a*_ *+ σ* ^*2*^_*e*_. To account for the binary nature of the trait i.e. 0 (mortality) or 1 (survival), heritability was adjusted to the underlying liability scale using the logit-link function which implies a correction of the residual variance by the factor π ^2^ /3. The genomic relationship matrix required for the analysis was obtained according to (VanRaden 2008) using the BLUPF90 software (Misztal *et al.* 2002) based on the final set of 17,919 filtered SNPs.

### Genome-wide association studies

Association analyses were run using the BLUPF90 software (Misztal *et al.* 2002) using a GBLUP approach and taking tank as a fixed effect. In this case, association analyses were performed for single SNPs and also for moving windows of 10, 20 and 50 adjacent SNPs (based on the linkage map position) that were created using POSTGSF90 (Aguilar *et al.* 2010).

### Genomic Selection

For the estimation of genomic prediction values, the genotype data used corresponded to the same 17,919 SNPs used for the GWAS analyses. Estimated breeding values were obtained using either pedigree-based BLUP (PBLUP) or Genomic best linear unbiased prediction (GBLUP) using the linear model described above. The accuracy of genomic selection was estimated by fivefold cross validation (training set 80%, validation set 20%), which were each repeated 10 times. Phenotypes (binary survival) from the validation population were masked and breeding values were estimated using ASReml 4 using the linear mixed model described above. Prediction accuracy was calculated as the correlation between the predicted EBVs of the validation set and the actual phenotypes divided by the square root of the heritability estimated in the validation population [∼r(y1, y2)/h]. Mean prediction accuracy values obtained from the different sets were compared between the pedigree and genomic approaches.

To assess the utility of low density SNP panels for breeding value prediction, two strategies for *in silico* selection of the SNPs were used. First, the low density SNP panel for use in the computing the genomic relationship matrix was selected by a progressive increase of the MAF threshold from 0.01 to 0.475 resulting in a progressive reduction in number of markers; Secondly, the low density SNP panel was selected using a strategy of random “thinning” of SNPs from the full dataset (15K, 10K, 5K, 2.5K, 1K, and down to 500 SNPs).

## Results

### Challenge survival and heritability

After the seven days of the disease challenge the average mortality per family for the entire challenge was 16.8 % and 24.7% for the LV and HV conditions, respectively, but with substantial variation in mortality levels between families (Table S2). High phenotypic correlations were observed between the family-level mortality rates in these two scenarios (R = 0.8; r^2^ = 0.64) as can be observed in (Figure 1 and Figure S1). Within the 718 genotyped samples (progenies) that were part of the challenge, the mortality level was 33 %.

**Figure 1.**
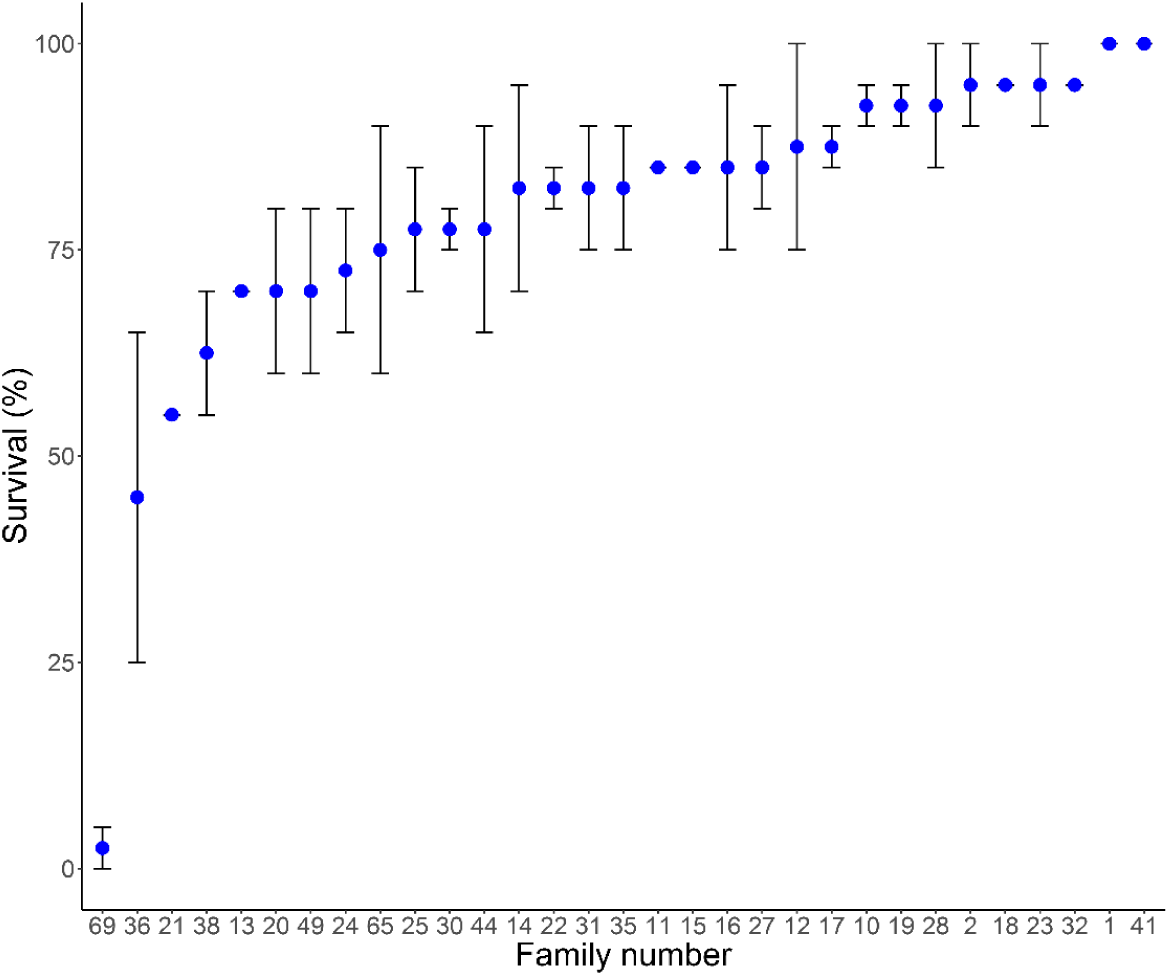
Average survival (s.e) for the 31 families included in the disease challenge

There were 762 individuals (718 progenies + 44 parents) with genotype data that passed QC and had accurate pedigree as confirmed by the family assignment software. Based on these data 30 families were consistent with the expected pedigree (although a small number of erroneously assigned individuals were identified and corrected based on genotype information. Only one family (id = 24) could not be assigned to both parents, suggesting that the dam was not included within the genotyped candidate parents, probably due to miss-labelling during the crosses. Making use of the updated pedigree information, the estimated heritability for the binary survival trait was 0.25 ± 0.05. These estimates were higher when using the genomic kinship matrix, with 0.37 ± 0.05. This gives evidence that there is a significant genetic component involved in the resistance to OsHV in Pacific oyster in these populations.

### Linkage map

The linkage mapping was performed using 30 full sibling families comprising 691 progenies and 43 parents (one family was discarded as both parents were not assigned). The linkage map contains 19,926 SNPs distributed on 10 LGs (in accordance with the Pacific oyster karyotype), with a length of 941 cM for the male map and 992 cM for the female map. 18,554 SNPs were mapped on the analysed families while the position of 1,372 SNPs was estimated from our previously published linkage map. Additionally, 14,058 SNPs are shared between the current and the previously published map obtained from a different population (Gutierrez *et al.* 2018a). The ∼20 K mapped SNPs correspond to 1,880 scaffolds and 133 contigs, according to the latest oyster genome assembly (GCA_000297895.1, Zhang *et al.* 2012). We observed that approximately 38 % (719 out 1,880) of the scaffolds with informative markers show evidence of errors in the assembly, due to assignment to at least two distinct LGs in our map, following similar pattern described in our previous linkage map (Gutierrez *et al.* 2018a; Hedgecock *et al.* 2015).

### Association analyses

GWAS performed using the BLUPF90 software did not detect any single SNPs showing a major association with the trait, shown as % of genetic variance explained by the SNP (Figure S2A). A similar situation was observed when a floating SNP window approach was taken, where variance values increased but not showing evidence of a major QTL (Figure 2 & Figure S2B-C). Particularly, regions on LG1, LG7, LG8 and LG10 seem to explain higher percentage of the genetic variance (between 1.2 and 1.67) although not high enough to suggest major-effect QTL, which implies a polygenic architecture to host resistance.

**Figure 2.**
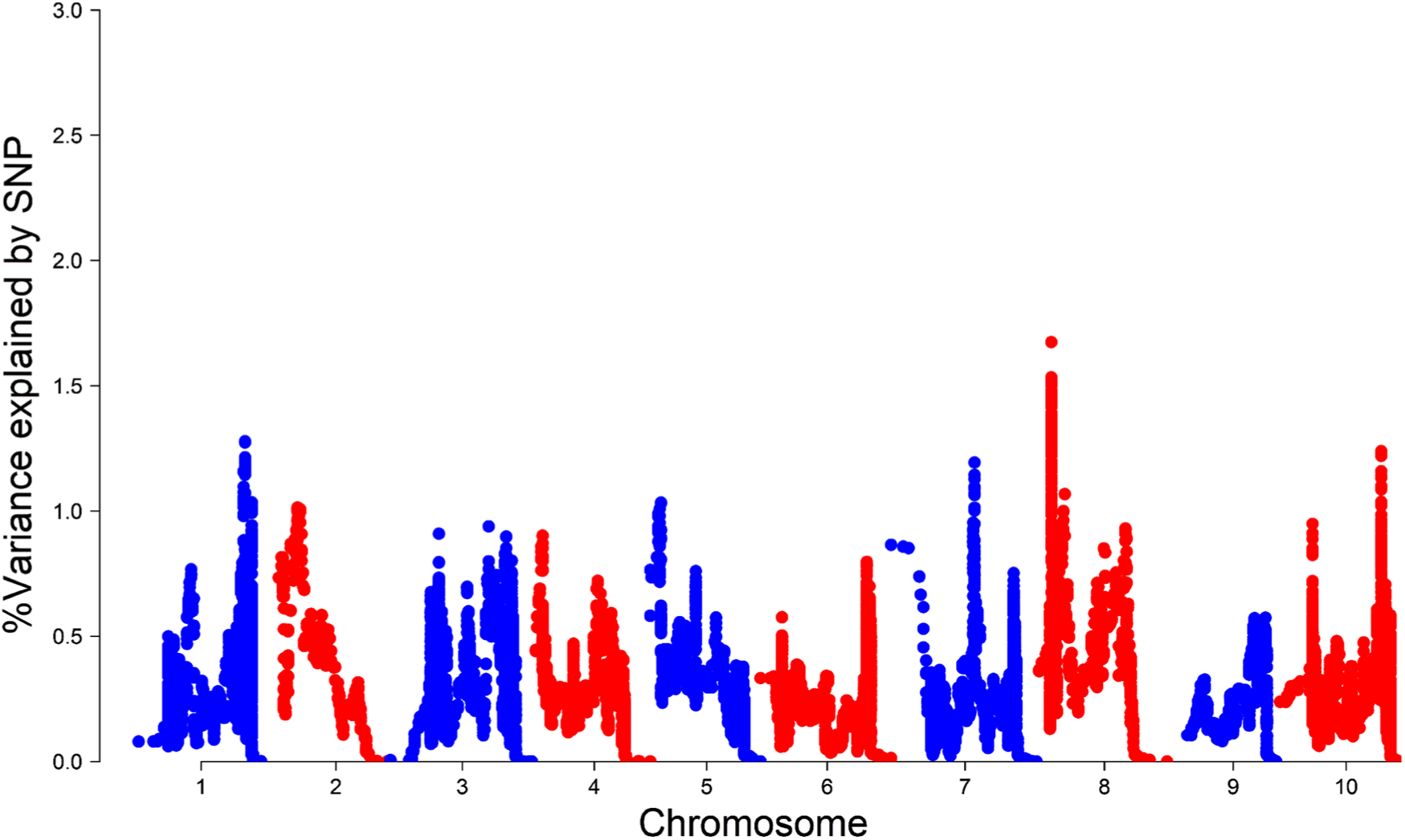
GWAS performed using moving windows of 50 SNPs, according to position on the linkage map.

### Genomic Prediction

Genomic prediction accuracy for the binary survival trait was analysed by randomly splitting the samples into training (80 %) and validation (20 %) sets for cross-validation, and this was repeated ten times. The genomic prediction accuracy results show that prediction accuracies obtained using genomic information (G-matrix) are higher than using the pedigree information (A-matrix) (Figure 3 & Table S3), with values ranging from 0.637 using pedigree to 0.758 using genomic information (increase of ∼19 %). Two approaches were taken to evaluate the effect of marker density on genomic prediction accuracy. The first used progressive increase of minor allele frequency (MAF) threshold, resulting in progressive decrease in SNP number. The second involved choosing subsets of SNPs for the low density panels at random. Both thinning approaches showed little impact on prediction accuracy until marker densities dropped below ∼2,500 SNPs. With the MAF approach, the genomic prediction accuracies obtained using the lower density SNP panels ranged from 0.755 to 0.693 (MAF>0.475 530 SNPs), while using the random subsets accuracies ranged from 0.758 to 0.678 (500 SNPs).

**Figure 3.**
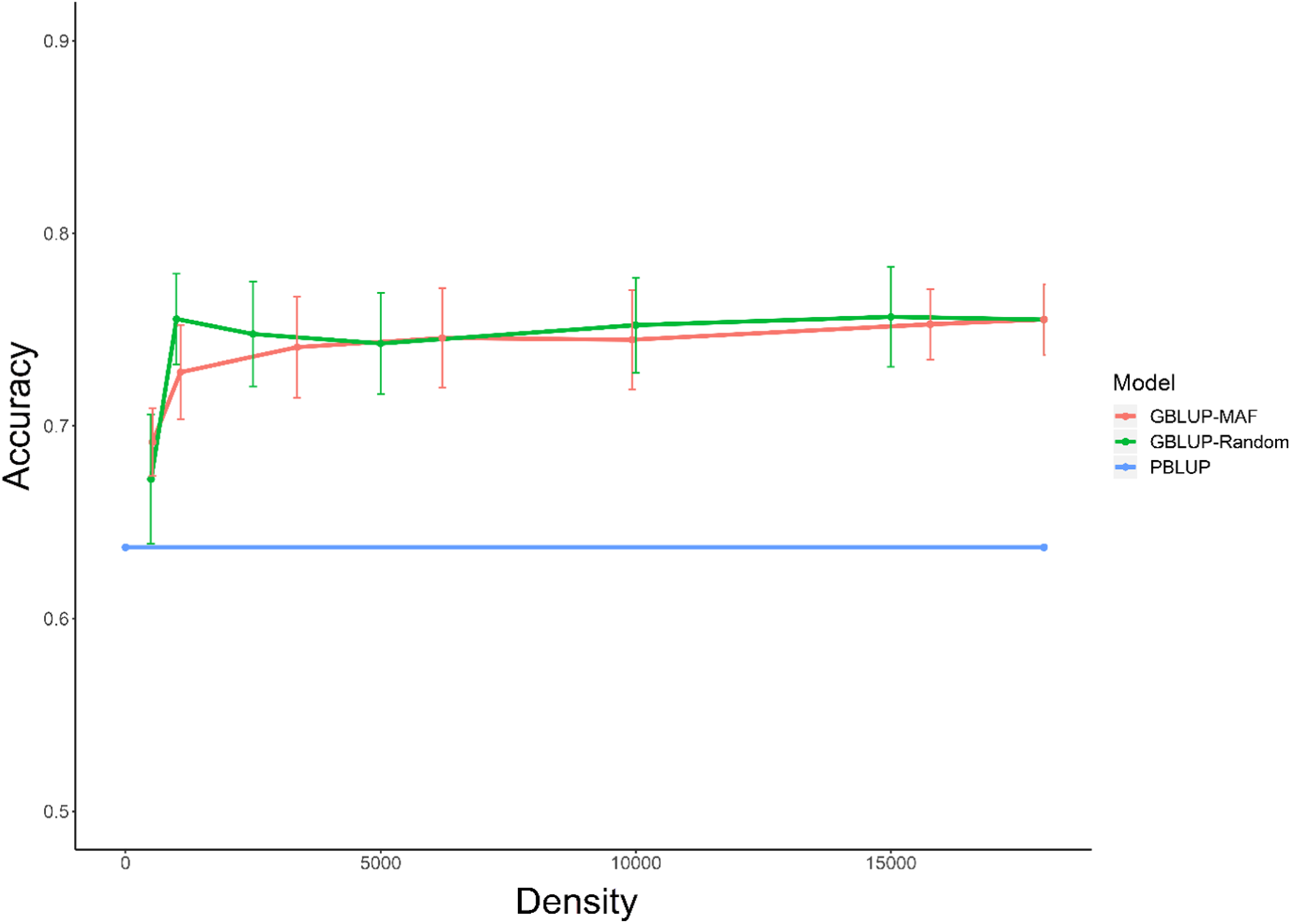
Genomic prediction results showing mean accuracies (s.d) obtained from both pedigree A-matrix (PBLUP) and genomic matrix G-matrix (GBLUP)

## Discussion

### Disease challenge and heritability of OsHV-1 resistance

The disease challenge set-up was consistent for both the low and high virus conditions showing high phenotypic correlation between family level mortality in the two challenges. This suggests that the variation in family level mortality is likely to be genetic, and the absence of mortality in the control group suggests that the challenge group mortality is due to OsHV-1 (Table S2). Moderate levels of heritability of OsHV resistance were observed in this study (0.25 - 0.37), which is similar to was has been described in previous studies (Dégremont *et al.* 2015b; Azéma *et al.* 2017; Camara *et al.* 2017), and higher than showed in our previous study in a different population with lower overall mortality rate (Gutierrez *et al.* 2018a). Heritability estimates were higher when using the genomic relationship matrix (GRM) compared to estimates obtained using the pedigree-based relationship matrix. A possible reason for this is overestimation of the additive genetic variance using a GRM due to high levels of linkage disequilibrium generated by recent selective breeding (Palaiokostas *et al.* 2018a; Fernando *et al.* 2017).

### Genome-wide association study

The linkage map construction resulted in ∼20K SNPs distributed across 10 linkage groups in similar positions as previously described in a different population (Gutierrez *et al.* 2018a). As with previous linkage maps, the mapping of SNPs located within single reference genome contigs to multiple linkage groups highlighted putative reference genome assembly errors (Hedgecock *et al.* 2015; Zhang *et al.* 2012). It is worth noting that 99.8% of the markers were located on the same LG as positioned in our previous map, which highlights the utility of the SNP chip across multiple populations, and the reliability of both maps.

The association analyses for OsHV-1 survival suggest that OsHV resistance in the Pacific oyster is likely controlled by multiple genomic regions in this population. Both the single SNP and moving window approach did not show evidence of a major region involved in the resistance. When the SNP window size was 50 the variance explained by some regions located on LG1, LG7, LG8 and LG10 was higher but no region explained more than 2 % of the genetic variance in the trait (Figure 2). Previous research into the genetics of OsHV resistance has also suggested an oligogenic or polygenic architecture of the trait which is consistent with our findings. In particular, a previous study found that LG V, VI, VII & IX (which correspond to LG 6, LG 7, LG 8 & LG 10 in our map) contain genomic regions associated to summer mortality resistance (Sauvage *et al.* 2010). Additionally, we recently described that regions on multiple LGs are likely involved in the resistance to OsHV in a Pacific oyster population from Guernsey (Gutierrez *et al.* 2018a). When taken together, these studies suggest that host resistance to OsHV in Pacific oyster is consistently a polygenic trait..

### Genomic selection

Genomic predictions (GBLUP) of breeding values for host resistance to OsHV are likely to be more accurate than those based on pedigree (PBLUP) in this population. The prediction accuracy values ranged from 0.678 to 0.758 for GBLUP (with SNP densities ranging from 500 to 18K), while PBLUP only reached an accuracy of 0.637. This result has been mirrored in other studies of genomic versus pedigree-based prediction of disease resistance breeding values for other important farmed fish species, e.g Atlantic salmon (Tsai *et al.* 2015; Yoshida *et al.* 2017; Ødegård *et al.* 2014; Robledo *et al.* 2018), rainbow trout (Vallejo *et al.* 2017; Yoshida *et al.* 2018), sea bream (Palaiokostas *et al.* 2016) and sea bass (Palaiokostas *et al.* 2018a). Further, in shellfish similar findings have been observed for prediction of breeding values for growth traits in scallop (Dou *et al.* 2016) and Pacific oyster (Gutierrez *et al.* 2018b). Therefore, the technical potential of genomic selection for expedited genetic improvement in shellfish has been shown. However, the economic viability remains an open question. The cost of genotyping is a key consideration for the commercial implementation of genomic selection in shellfish. In the current study, <1000 SNPs was a sufficient density to provide ∼19 % increase in prediction accuracy versus PBLUP. This raises the possibility of a low density genomic selection approach for OsHV-1 resistance in oyster breeding, since low density genotyping can be substantially cheaper than high density SNP arrays. It is worth noting that the training and validation sets in the current study contain closely related animals, including full siblings. As a result, these individuals will share large genomic segments, which can be capture by the low density SNP panels. As the genetic distance between the training and validation sets increased, genomic prediction accuracy is likely to decrease markedly, as has been shown in other aquaculture species (Palaiokostas et al., 2019). In addition, it has been demonstrated some populations of Pacific oysters exhibit rapid decay of linkage disequilibrium (Gutierrez et al. 2017). As such, regular testing on close relatives of breeding candidates is required to maintain prediction accuracy, otherwise accuracy will decrease with successive generations of a breeding programme. Having that said, this scenario is typical for aquaculture breeding programs where testing of full-siblings of selection candidates is performed. Historically, the breeding programme for Pacific oysters run by the Cawthron Institute has used broodstock which are put through grow-out in field environments and are brought back into the hatchery for reproduction. As such, when OsHV outbreaks occur, it has been common practice to breed from survivors of the highest performing families. Genomic selection is best-suited to traits that are not measurable on the selection candidates themselves, and helps selection of individuals from within a full sibling family. When survivors are used as breeding candidates, these benefits are somewhat negated. However, in the event that broodstock cannot be brought back into a hatchery (e.g. due to a biosecurity issue) the practical value of genomic selection would be much higher.

The Cawthron breeding programme found no evidence for the vertical transmission of OsHV-1 in the hatchery, and the existing presence of OsHV-1 in wild populations meant that the biosecurity implications of breeding from survivors were manageable. Therefore the ability to select broodstock from large on-farm progeny trials exposed to OsHV-1 enabled the benefits of within-family selection to be captured, rather than relying only on among-family selection. Where vertical transmission occurs in the hatchery or biosecurity is constraining, the option to breed from survivors may not be available. In this case, genomic selection provides the opportunity for within family selection even though the phenotype (e.g. survival) cannot be measured directly on broodstock candidates.

## Conclusion

The results from the current study provide evidence indicating that OsHV resistance is polygenic in Pacific oyster, consistent with current literature and analyses across different populations. Due to the polygenic nature of the resistance, genomic selection is a well-placed methodology for the improvement of current pedigree-based selection schemes. Indeed results show that genomic prediction of OsHV-1 resistance is more accurate than pedigree-based prediction even with a reduced number of SNPs (down to less than 1,000 SNPs). These results suggest that low cost genotyping solutions could be within reach to provide a G-matrix capable of generating accurate GEBV values. The use of genomic data is likely to bring significant improvement to Pacific oyster breeding programmes, particularly to improve selection of challenging traits that rely on sib-testing (e.g. disease resistance) where genomic selection can more effectively capture within-family variation.

## Acknowledgements

The authors gratefully acknowledge funding from BBSRC and NERC under the United Kingdom Aquaculture Initiative (BB/M026140/1 and NE/P010695/1) in addition to BBSRC Institute Strategic Program Grants (BBS/E/D/20002172 and BBS/E/D/30002275). Edinburgh Genomics was partly supported through core grants from NERC (R8/H10/56), MRC (MR/K001744/1), and BBSRC (BB/J004243/1). Cawthron Institute was funded by New Zealand’s MBIE funded Cultured Shellfish programme (CAWX1315).

## References

Aguilar, I., I. Misztal, D.L. Johnson, A. Legarra, S. Tsuruta et al., 2010 Hot topic: A unified approach to utilize phenotypic, full pedigree, and genomic information for genetic evaluation of Holstein final score. J. Dairy Sci. 93 (2):743–752.

Aulchenko, Y.S., S. Ripke, A. Isaacs, and C.M. van Duijn, 2007 GenABEL: an R library for genome-wide association analysis. Bioinformatics 23 (10):1294–1296.

Azéma, P., J.-B. Lamy, P. Boudry, T. Renault, M.-A. Travers et al., 2017 Genetic parameters of resistance to Vibrio aestuarianus, and OsHV-1 infections in the Pacific oyster, Crassostrea gigas, at three different life stages. Genetics Selection Evolution 49 (1):23.

Barría, A., K.A. Christensen, G.M. Yoshida, K. Correa, A. Jedlicki et al., 2018 Genomic Predictions and Genome-Wide Association Study of Resistance Against *Piscirickettsia salmonis* in Coho Salmon (*Oncorhynchus kisutch*) Using ddRAD Sequencing. G3: Genes|Genomes|Genetics 8 (4):1183–1194.

Camara, M.D., and J.E. Symonds, 2014 Genetic improvement of New Zealand aquaculture species: programmes, progress and prospects. N. Z. J. Mar. Freshwater Res. 48 (3):466–491.

Camara, M.D., S. Yen, H.F. Kaspar, A. Kesarcodi-Watson, N. King et al., 2017 Assessment of heat shock and laboratory virus challenges to selectively breed for ostreid herpesvirus 1 (OsHV-1) resistance in the Pacific oyster, Crassostrea gigas. Aquaculture 469:50–58.

Chang, C.C., C.C. Chow, L.C. Tellier, S. Vattikuti, S.M. Purcell et al., 2015 Second-generation PLINK: rising to the challenge of larger and richer datasets. GigaScience 4 (1):7.

de Lorgeril, J., A. Lucasson, B. Petton, E. Toulza, C. Montagnani et al., 2018 Immune-suppression by OsHV-1 viral infection causes fatal bacteraemia in Pacific oysters. Nature Communications 9 (1):4215.

Dégremont, L., E. Bédier, and P. Boudry, 2010 Summer mortality of hatchery-produced Pacific oyster spat (Crassostrea gigas). II. Response to selection for survival and its influence on growth and yield. Aquaculture 299 (1):21–29.

Dégremont, L., C. Garcia, and S.K. Allen, 2015a Genetic improvement for disease resistance in oysters: A review. J. Invertebr. Pathol. 131:226–241.

Dégremont, L., J.-B. Lamy, J.-F. Pépin, M.-A. Travers, and T. Renault, 2015b New Insight for the Genetic Evaluation of Resistance to Ostreid Herpesvirus Infection, a Worldwide Disease, in *Crassostrea gigas*. PLoS One 10 (6):e0127917.

Dégremont, L., M. Nourry, and E. Maurouard, 2015c Mass selection for survival and resistance to OsHV-1 infection in Crassostrea gigas spat in field conditions: response to selection after four generations. Aquaculture 446:111–121.

Dou, J., X. Li, Q. Fu, W. Jiao, Y. Li et al., 2016 Evaluation of the 2b-RAD method for genomic selection in scallop breeding. Sci. Rep. 6:19244.

Fernando, R.L., H. Cheng, X. Sun, and D.J. Garrick, 2017 A comparison of identity-by-descent and identity-by-state matrices that are used for genetic evaluation and estimation of variance components. Journal of Animal Breeding and Genetics 134 (3):213–223.

Gutierrez, A.P., T.P. Bean, C. Hooper, C.A. Stenton, M.B. Sanders et al., 2018a A Genome-Wide Association Study for Host Resistance to Ostreid Herpesvirus in Pacific Oysters (*Crassostrea gigas*). G3: Genes|Genomes|Genetics 8 (4):1273–1280.

Gutierrez, A.P., O. Matika, T.P. Bean, and R.D. Houston, 2018b Genomic selection for growth traits in Pacific oyster (Crassostrea gigas): Potential of low-density marker panels for breeding value prediction. Frontiers in Genetics 9:391.

Gutierrez, A.P., F. Turner, K. Gharbi, R. Talbot, N.R. Lowe et al., 2017 Development of a Medium Density Combined-Species SNP Array for Pacific and European Oysters (*Crassostrea gigas* and *Ostrea edulis*). G3: Genes|Genomes|Genetics 7 (7):2209–2218.

Hedgecock, D., G. Shin, A.Y. Gracey, D.V. Den Berg, and M.P. Samanta, 2015 Second-Generation Linkage Maps for the Pacific Oyster Crassostrea gigas Reveal Errors in Assembly of Genome Scaffolds. G3: Genes|Genomes|Genetics 5 (10):2007–2019.

Houston, R.D., 2017 Future directions in breeding for disease resistance in aquaculture species. Revista Brasileira de Zootecnia 46:545–551.

Houston, R.D., J.B. Taggart, T. Cézard, M. Bekaert, N.R. Lowe et al., 2014 Development and validation of a high density SNP genotyping array for Atlantic salmon (Salmo salar). BMC Genomics 15 (1):90.

Kalinowski, S.T., M.L. Taper, and T.C. Marshall, 2007 Revising how the computer program cervus accommodates genotyping error increases success in paternity assignment. Mol. Ecol. 16 (5):1099–1106.

Liu, S., L. Sun, Y. Li, F. Sun, Y. Jiang et al., 2014 Development of the catfish 250K SNP array for genome-wide association studies. BMC Res. Notes 7 (1):135.

Malham, S.K., E. Cotter, S. O’Keeffe, S. Lynch, S.C. Culloty et al., 2009 Summer mortality of the Pacific oyster, Crassostrea gigas, in the Irish Sea: The influence of temperature and nutrients on health and survival. Aquaculture 287 (1):128–138.

Meuwissen, T.H.E., B.J. Hayes, and M.E. Goddard, 2001 Prediction of Total Genetic Value Using Genome-Wide Dense Marker Maps. Genetics 157 (4):1819–1829.

Misztal, I., S. Tsuruta, T. Strabel, B. Auvray, T. Druet et al., 2002 BLUPF90 and related programs (BGF90), pp. 743–744 in Proceedings of the 7th world congress on genetics applied to livestock production.

Ødegård, J., T. Moen, N. Santi, S.A. Korsvoll, S. Kjøglum et al., 2014 Genomic prediction in an admixed population of Atlantic salmon (Salmo salar). Frontiers in Genetics 5 (402).

Palaiokostas C., T. Vesely, M. Kocour, M. Prchal, D. Pokorova, V. Piackova, L. Pojezdal, R.D. Houston. 2014 Optimizing Genomic Prediction of Host Resistance to Koi Herpesvirus Disease in Carp. Frontiers in Genetics 10(543).

Palaiokostas, C., S. Cariou, A. Bestin, J.-S. Bruant, P. Haffray et al., 2018a Genome-wide association and genomic prediction of resistance to viral nervous necrosis in European sea bass (Dicentrarchus labrax) using RAD sequencing. Genetics Selection Evolution 50 (1):30.

Palaiokostas, C., S. Ferraresso, R. Franch, R.D. Houston, and L. Bargelloni, 2016 Genomic Prediction of Resistance to Pasteurellosis in Gilthead Sea Bream (*Sparus aurata*) Using 2b-RAD Sequencing. G3: Genes|Genomes|Genetics 6 (11):3693–3700.

Palaiokostas, C., M. Kocour, M. Prchal, and R.D. Houston, 2018b Accuracy of Genomic Evaluations of Juvenile Growth Rate in Common Carp (Cyprinus carpio) Using Genotyping by Sequencing. Frontiers in Genetics 9 (82).

Palti, Y., G. Gao, S. Liu, M.P. Kent, S. Lien et al., 2015 The development and characterization of a 57K single nucleotide polymorphism array for rainbow trout. Mol. Ecol. Resour. 15 (3):662–672.

Petton, B., M. Bruto, A. James, Y. Labreuche, M. Alunno-Bruscia et al., 2015 Crassostrea gigas mortality in France: the usual suspect, a herpes virus, may not be the killer in this polymicrobial opportunistic disease. Front. Microbiol. 6 (686).

Qi, H., K. Song, C. Li, W. Wang, B. Li et al., 2017 Construction and evaluation of a high-density SNP array for the Pacific oyster (Crassostrea gigas). PLoS One 12 (3):e0174007.

Rastas, P., 2017 Lep-MAP3: robust linkage mapping even for low-coverage whole genome sequencing data. Bioinformatics.

Robledo, D., O. Matika, A. Hamilton, and R.D. Houston, 2018 Genome-Wide Association and Genomic Selection for Resistance to Amoebic Gill Disease in Atlantic Salmon. G3: Genes|Genomes|Genetics 8 (4):1195–1203.

Sauvage, C., P. Boudry, D.J. De Koning, C.S. Haley, S. Heurtebise et al., 2010 QTL for resistance to summer mortality and OsHV-1 load in the Pacific oyster (Crassostrea gigas). Anim. Genet. 41 (4):390–399.

Segarra, A., J.F. Pépin, I. Arzul, B. Morga, N. Faury et al., 2010 Detection and description of a particular Ostreid herpesvirus 1 genotype associated with massive mortality outbreaks of Pacific oysters, Crassostrea gigas, in France in 2008. Virus Res. 153 (1):92–99.

Tsai, H.-Y., A. Hamilton, A.E. Tinch, D.R. Guy, K. Gharbi et al., 2015 Genome wide association and genomic prediction for growth traits in juvenile farmed Atlantic salmon using a high density SNP array. BMC Genomics 16 (1):969.

Vallejo, R.L., T.D. Leeds, G. Gao, J.E. Parsons, K.E. Martin et al., 2017 Genomic selection models double the accuracy of predicted breeding values for bacterial cold water disease resistance compared to a traditional pedigree-based model in rainbow trout aquaculture. Genetics Selection Evolution 49 (1):17.

Vallejo, R.L., R.M.O. Silva, J.P. Evenhuis, G. Gao, S. Liu et al., 2018 Accurate genomic predictions for BCWD resistance in rainbow trout are achieved using low-density SNP panels: Evidence that long-range LD is a major contributing factor. Journal of Animal Breeding and Genetics 135 (4):263–274.

VanRaden, P.M., 2008 Efficient Methods to Compute Genomic Predictions. J. Dairy Sci. 91 (11):4414–4423.

Xu, J., Z. Zhao, X. Zhang, X. Zheng, J. Li et al., 2014 Development and evaluation of the first high-throughput SNP array for common carp (Cyprinus carpio). BMC Genomics 15 (1):307.

Yáñez, J.M., S. Naswa, M.E. López, L. Bassini, K. Correa et al., 2016 Genomewide single nucleotide polymorphism discovery in Atlantic salmon (Salmo salar): validation in wild and farmed American and European populations. Mol. Ecol. Resour. 16 (4):1002–1011.

Yoshida, G.M., R. Bangera, R. Carvalheiro, K. Correa, R. Figueroa et al., 2018 Genomic Prediction Accuracy for Resistance Against *Piscirickettsia salmonis* in Farmed Rainbow Trout. G3: Genes|Genomes|Genetics 8 (2):719–726.

Yoshida, G.M., J.P. Lhorente, R. Carvalheiro, and J.M. Yáñez, 2017 Bayesian genome-wide association analysis for body weight in farmed Atlantic salmon (Salmo salar L.). Anim. Genet. 48 (6):698–703.

Zhang, G., X. Fang, X. Guo, L. Li, R. Luo et al., 2012 The oyster genome reveals stress adaptation and complexity of shell formation. Nature 490 (7418):49–54.

