## Supplementary material for "Potential of genomic selection for improvement of resistance to Ostreid Herpes virus in Pacific oyster (*Crassostrea gigas*)": Figure S1 & S2

Figure S1. Correlation of survival level observed per family in both the low virus (LV) and high virus (HV) conditions

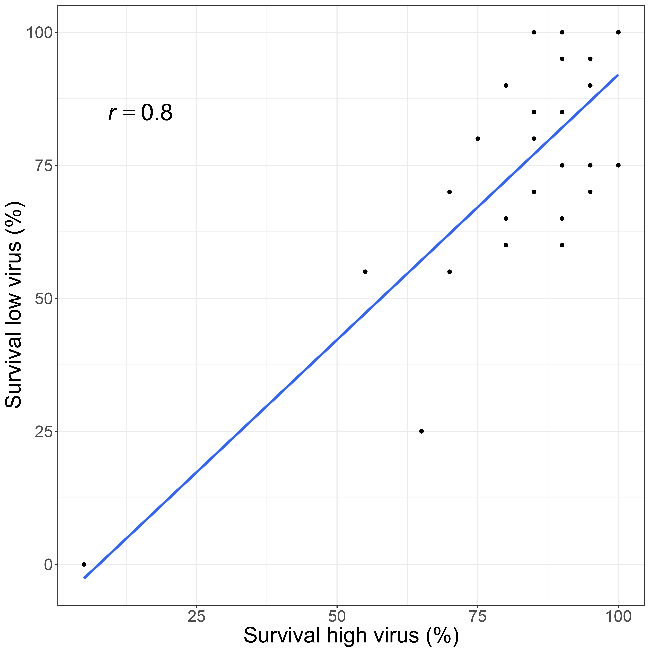

Figure S2. GWAS results for OsHV survival based on: A. single SNP; B. 10 SNP window; C. 20 SNP window.

**A**

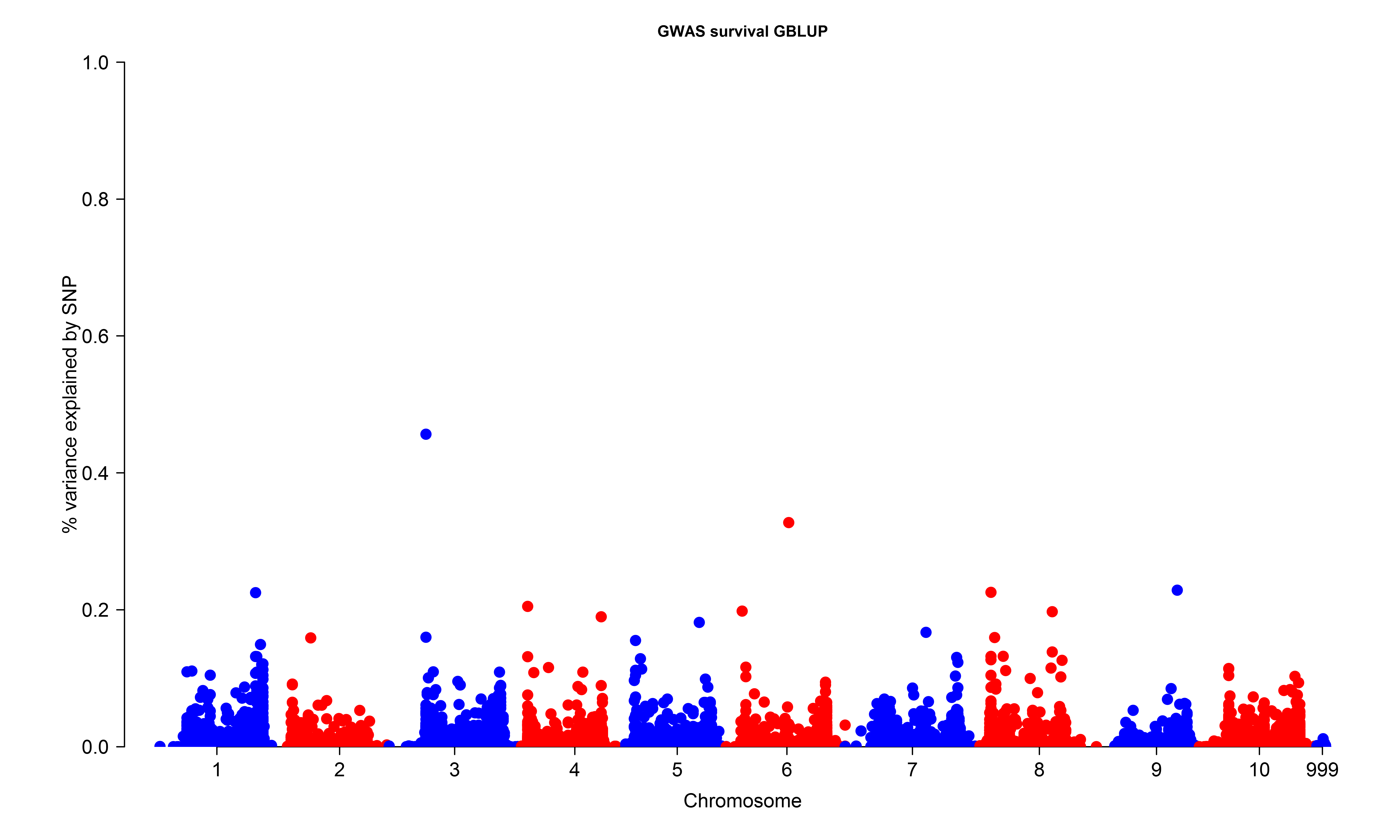

**B**

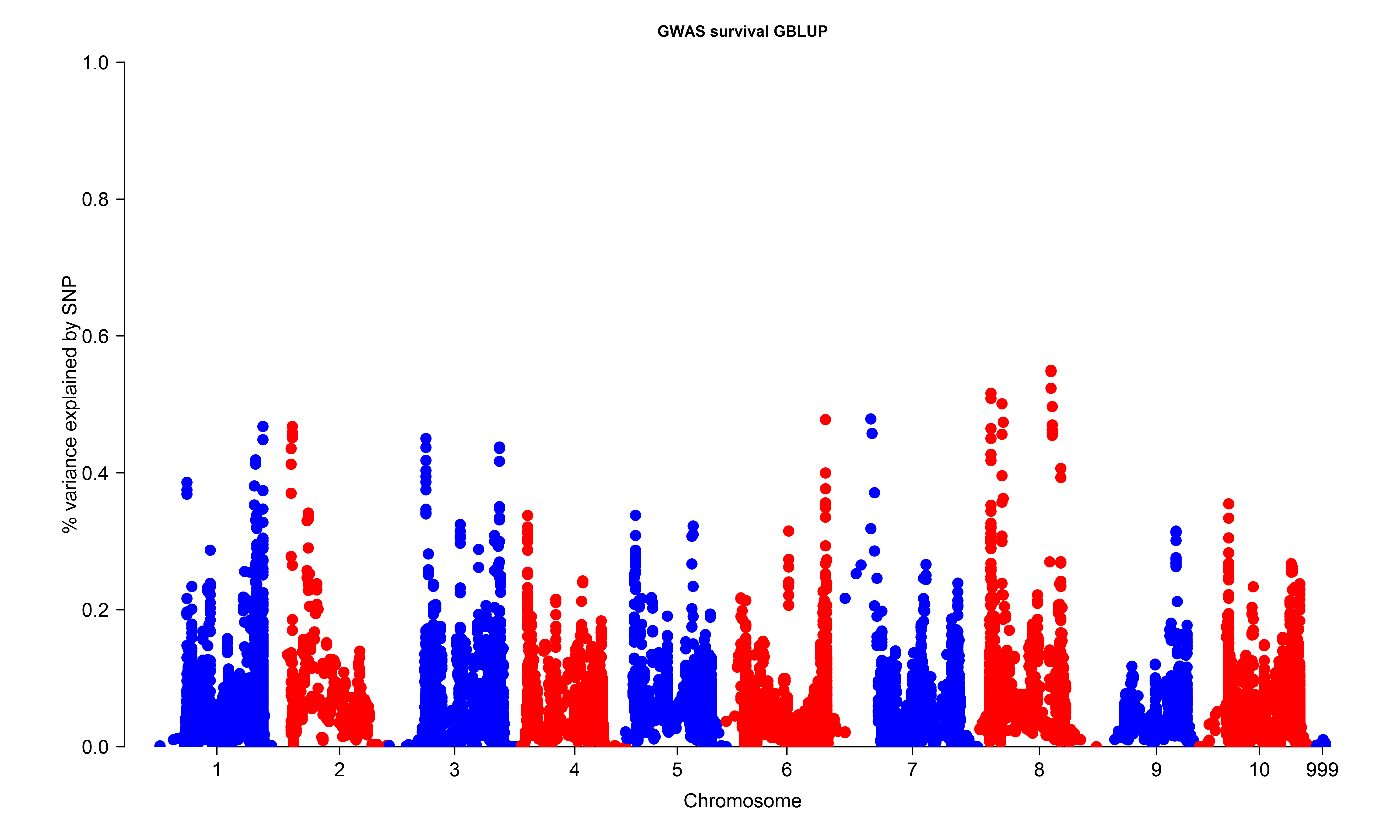

**C**

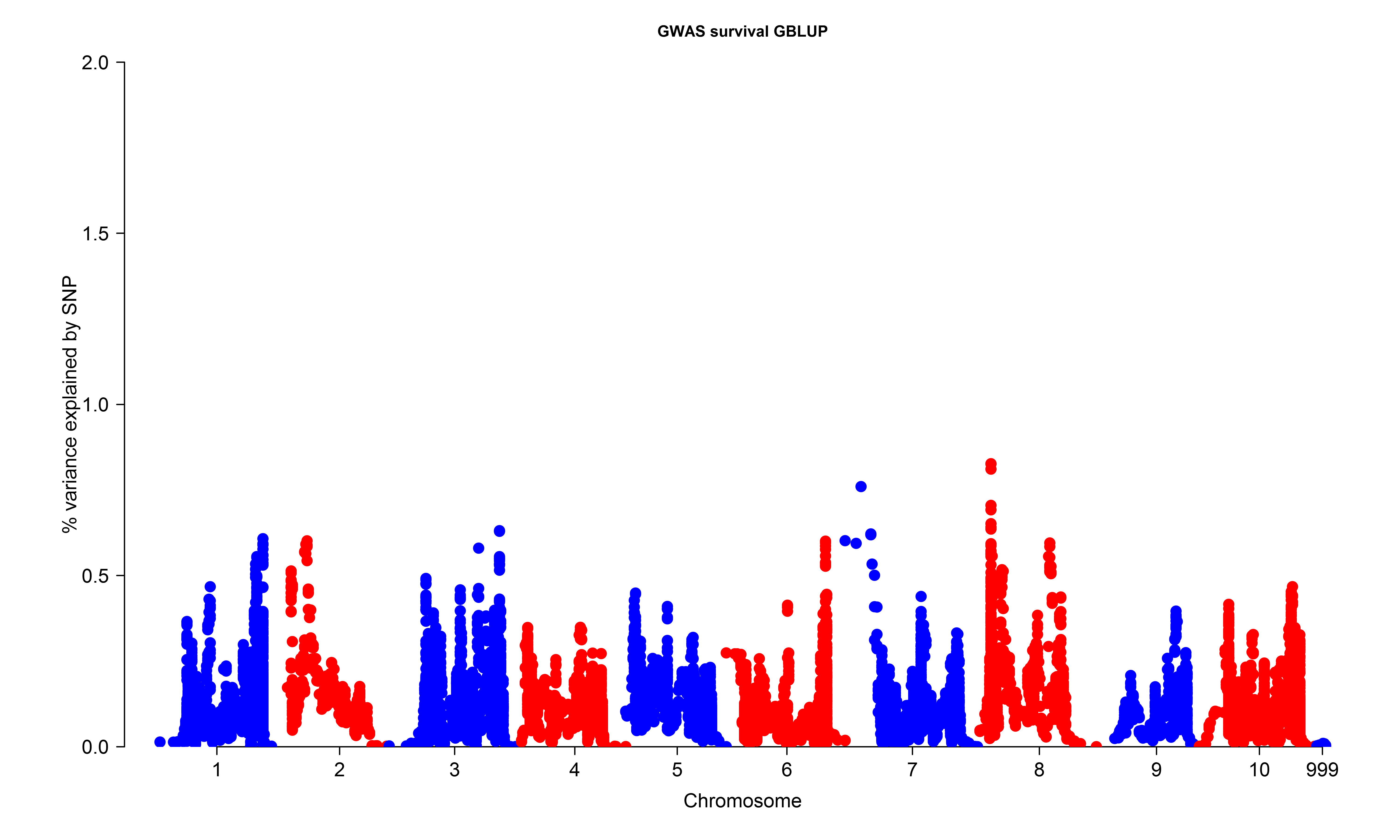
